# Single-cell ID-seq identifies BMP signaling as a driver of a late stage epidermal differentiation program

**DOI:** 10.1101/350850

**Authors:** Roderick E. van Eijl, Jessie A.G. van Buggenum, Sabine E.J. Tanis, Joost Hendriks, Klaas W. Mulder

## Abstract

Epidermal homeostasis requires balanced and coordinated adult stem cell renewal and differentiation. These processes are controlled by both extracellular signaling and by cell intrinsic transcription regulatory networks, yet how these control mechanisms are integrated to achieve this unclear. Here, we developed single-cell ID-seq and measured 69 antibody-DNA conjugates (including 34 phospho-specific epitopes) to study the activation state of signaling pathways during epidermal differentiation at the single-cell level. Computational pseudo-timing inference revealed activation of the JAK-STAT, WNT and BMP pathways along the epidermal differentiation trajectory. During differentiation, cells start producing BMP2 ligands and activate the canonical intracellular effectors SMAD1/5/9. Mechanistically, the BMP pathway is responsible for directly activating a specific transcription program that includes the key differentiation transcription factors MAF and MAFB to allow terminal differentiation. We propose that incorporating autocrine signaling pathway activation into a transcription regulatory network enables regional coordination of transcription programs during epidermal differentiation.

## Introduction

The human epidermis is continuously turned-over throughout life, a process that requires control and coordination of stem cell renewal and differentiation. During epidermal homeostasis, proliferating stem/progenitor cells residing in the basal layer replenish terminally differentiated cells that are shed from the skin surface (Blanpain and Fuchs, 2006; Solanas and Benitah, 2013; Watt et al., 2006). Human epidermal stem cells can be maintained in culture and used to regenerate functional epidermis *in vivo* upon transplantation and retain their capacity to differentiate *in vitro* (Barrandon et al., 2012; Green, 2008; Hirsch et al., 2017; Rheinwald and Green, 1975). The process of differentiation is driven forward by consecutive activation of transcriptional programs, yet the mechanisms underlying their sequence and timing are not well understood. In the epidermal basal layer, cells receive proliferative signals, predominantly via the epidermal growth factor receptor, and contact the underlying basement membrane (Watt, 2002; Watt and Huck, 2013). These contacts are mediated by focal adhesions and hemi-desmosomes that contain integrin beta-1 and integrin alpha-6, respectively (Watt, 2002; Watt and Huck, 2013). As cells stop proliferating and initiate differentiation, these adhesion structures are resolved, allowing the cells to detach from the basement membrane and migrate up towards the epidermal surface. During this migration, the cells undergo major transitions in transcriptional programs, eventually producing the terminally differentiated keratinocytes that form the outermost protective layer of our skin. TP63 is a key DNA-binding transcription factor in epidermal stem cell renewal and upon differentiation its expression is decreased (Kouwenhoven et al., 2010; Truong et al., 2006). In contrast, other transcription factors, including KLF4, OVOL2, GRHL3, MAF/MAFB and ZNF750, are induced (Bhaduri et al., 2015; Koster and Roop, 2004; Lopez-Pajares et al., 2015; Sen et al., 2012; Wells et al., 2009). Of these, MAF and MAFB cooperatively regulate a transcription program that includes ZNF750, which subsequently drives expression of terminal differentiation genes (Lopez-Pajares et al., 2015). This concept of sequential activation of transcription factors (also called transcription regulatory networks) explains cell intrinsic progression of epidermal differentiation. Indeed, human keratinocytes differentiate when placed in conditions where they are not in contact with other cells, for instance in suspension in methylcellulose or on micro-patterned islands, (Adams and Watt, 1989; Connelly et al., 2010). However, this does not take into account the need for regional coordination of differentiation in a tissue context. For instance, the basal, spinous, granular and cornified layers of the epidermis are morphologically distinct and can be distinguished using specific markers reflecting differences in transcriptional programs. The fact that these are sequentially formed layers indicates the need for a form of coordination that is not immediately explained by the function of cell intrinsic transcription factor networks.

Several signaling pathways (e.g. Integrins, EGF, TGFbeta and Notch) have been implicated in human epidermal differentiation, yet their temporal ordering and functional contribution to the control of specific transcription programs remains largely unknown (Beck and Blanpain, 2012; Blanpain and Fuchs, 2006; Li et al., 2003; Watt, 2002; Watt et al., 2006). These pathways depend on binding of a peptide ligand to the extracellular part of a transmembrane receptor. This receptor then relays this interaction into an intracellular cascade, usually involving multiple kinases and phosphorylation events to regulate specific transcription programs. As such, activation of extracellular signaling pathways may serve as a self-contained timing mechanism to drive differentiation forward in a tissue and safeguard the irreversibility of the process.

The activity of a signaling pathway can be monitored through the activated (phosphorylated) states of its components. However, populations of cells rarely display homogeneous timing of differentiation (Altschuler and Wu, 2010; Wu and Singh, 2012). We therefore set out to study signaling pathway activity during human epidermal cell differentiation at the single-cell level.

## Results

### scID-seq allows robust, reproducible and highly multiplexed protein detection in single cells

We recently described the Immuno-Detection by sequencing (ID-seq) technology to quantify >70 proteins and phospho-proteins in hundreds of cell populations simultaneously (Buggenum et al., 2017; Van Buggenum et al., 2016). In short, ID-seq entails immuno-staining with DNA-barcoded antibodies followed by quantification through high-throughput sequencing and barcode counting. We now adapted the ID-seq workflow to include immuno-staining of cells in suspension followed by flow cytometry-based single-cell distribution into 96-wells PCR plates to prepare samples for sequencing (figure 1A and figure S1). The inclusion of a 15 nucleotide unique molecular identifier (UMI) allowed counting-based quantification and ensured that potential duplication artifacts introduced during sample preparation can be accounted for (Kivioja et al., 2012). After sequencing, we found that counts from wells containing a single sorted cell were 100-fold higher than empty wells, suggesting that only 1% of the signal constituted technical background (figure 1B). Next, we sought to characterize the reproducibility/confidence of quantification of antibody-derived UMI-counts within single cells. For this, we generated a panel of five antibodies against cell surface (ITGA6 and ITGB1), cytoplasmic (Actin and TGM1) and nuclear (RNApol2) epitopes, each of which were independently conjugated to 9 distinct DNA-barcodes, generating a total pool of 45 antibody-DNA conjugates. Each of these barcodes serves as a technical replicate and their concordance therefore reflects the reproducibility of single cell ID-seq measurements. All of the barcodes for each of these five antibodies showed very good correlation across cells (R=0.95-0.99), indicating a low level of noise in scID-seq measurements (figure 1C and S2). To further validate scID-seq, we stained human epidermal stem cells simultaneously with separate ITGB1 antibodies containing a fluorescent group, or the DNA-barcode. We recorded the FACS-based ITGB1 signal, as well as the ITGB1 DNA-barcode counts for individual cells. This revealed that scID-seq counts indeed reflect standard FACS measurements for the same cell (R=0.76, figure 1D). Together, these results establish single-cell ID-seq as a robust and reproducible method to measure proteins in individual cells. Moreover, the fact that each DNA-barcode is unique to a specific antibody allows many antibodies to be multiplexed and measured in each individual cell.

**Figure 1:**
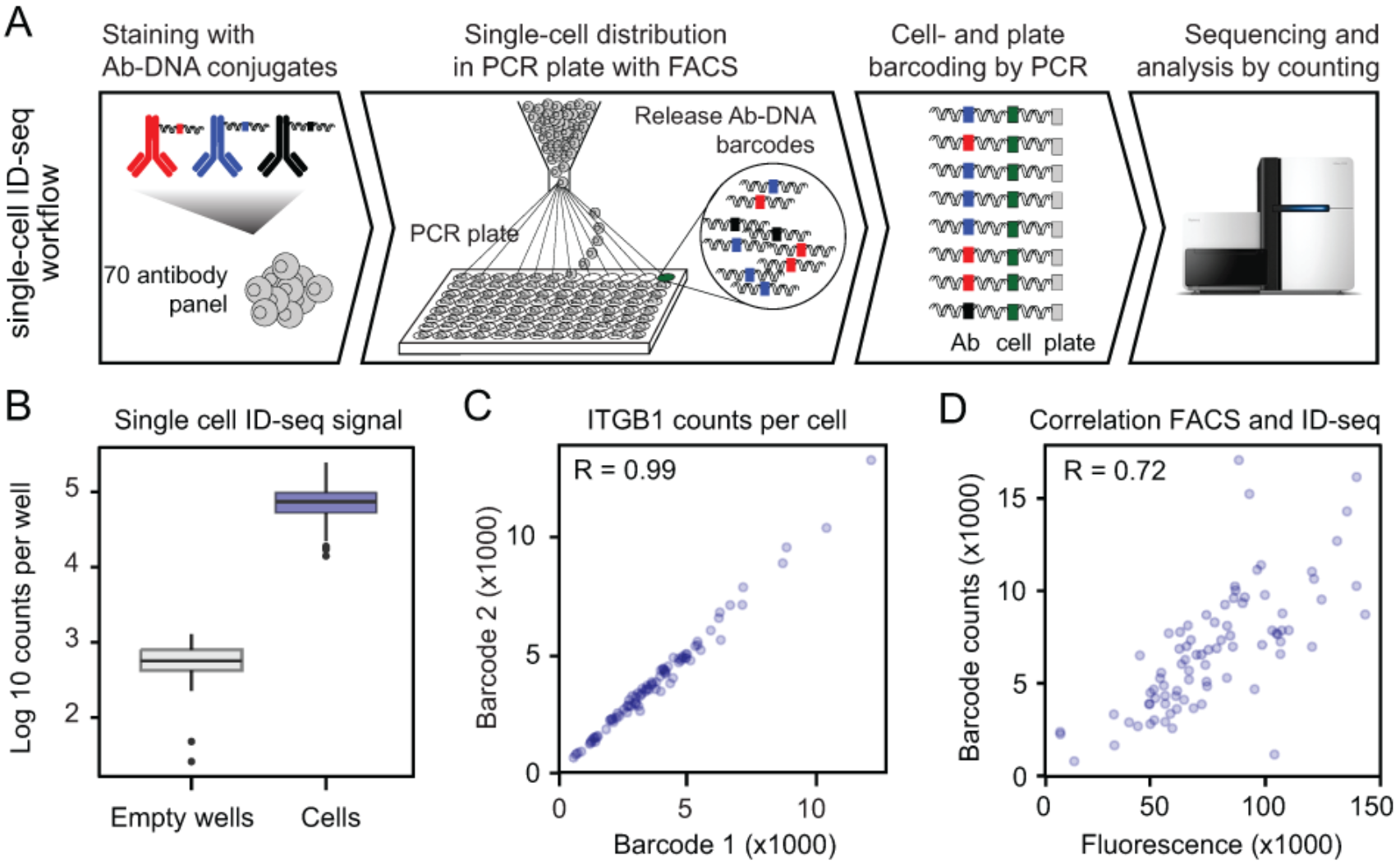
Single-cell Immuno-Detection by Sequencing (scID-seq) enables robust and reproducible protein detection in individual cells. **(A)** Schematic overview of the scID-seq workflow. **(B)** Single cell derived antibody counts indicate low technical noise levels in scID-seq. Single cells stained with antibody-DNA conjugates sorted into individual wells of a 96-well plate and compared to wells were no cells were sorted into using scID-seq (n=84 and 24, respectively). **(C)** scID-seq reproducibly measures antibody signals in single cells. Independently generated antibody-DNA conjugates were used to stain human epidermal stem cells. Barcode counts derived from the two barcodes were plotted against each other, indicating high reproducibility of the protein measurements from the same cell. **(D)** scID-seq reflects Fluorescent Activated Cell Sorting measurements. ITGB1 levels were measured in human epidermal stem cells using FACS and scID-seq. Fluorescent signal and scID-seq counts for each cell (n=84) showed a good correlation (R=0.76).

### scID-seq distinguishes renewing and differentiated epidermal cells

We applied single-cell ID-seq to monitor the activity of signaling pathways and other cellular processes during epidermal differentiation using a panel of 70 antibody-DNA conjugates (Buggenum et al., 2017; Van Buggenum et al., 2016). These antibodies cover a broad range of cellular processes including cell cycle, DNA damage, epidermal self-renewal and differentiation, as well as the intracellular signalling status for the EGF, G-protein coupled receptors, calcium signalling, TNFα, TGFβ, Notch, WNT and BMP pathways (Buggenum et al., 2017 and supplemental table 1). This panel includes 34 antibodies against phosphorylated epitopes and was previously validated and used to identify kinases involved in epidermal renewal (Buggenum et al., 2017). For 11 of the phosphorylated epitopes we were also able to measure the non-phosphorylated protein, allowing us to correct for total protein levels by calculating the phosphorylated to total protein ratio per cell. Most of the targeted processes are measured using multiple (3-5) independent validated antibodies (Buggenum et al., 2017).

The surface level of ITGB1, reflects a cell’s potential to self-renew (Watt, 2002). We used FACS isolation of single cells based on their ITGB1 levels to capture the transitions that underlie the differentiation process (figure 2A). Colony formation assays confirmed the loss of proliferative capacity in cells expressing low levels of ITGB1 (figure S3A). After staining with the 70 antibody-DNA conjugates, cells were sorted into ITGB1 positive and ITGB1low populations based on their fluorescent ITGB1 antibody signal and subjected to scID-seq. After quality control and filtering we obtained a dataset of 220 single cells in which 69 antibody-DNA conjugates were quantified. For each cell, antibody-DNA counts were normalized for differences in sequencing depth by subsampling. To avoid downstream analyses from being dominated by the most abundant epitopes, we scaled all antibody counts from 0 to 1 across all cells. Unsupervised principal component analysis separated the integrin beta-1 low (differentiated) cells from the ITGB1+ cells, confirming the notion that these represent distinct cell states (figure S3B). Although we isolated cells only with respect to their cell surface expression of ITGB1, we found that other proteins displayed concordant dynamics. For instance, the basal cell markers ITGA6 and TP63 were also decreased in the ITGB1low population (figure 2B). In contrast, this population exhibited higher expression of the differentiation-associated proteins TGM1, Notch1 Intracellular Domain (NICD) and KLF4 (figure 2B). Furthermore, cell cycle markers, such as Rb-p and cdc2 (reflecting the G1-S and G2-M transitions, respectively), distinguished these populations very well (figure 2B). This is consistent with the loss of proliferative capacity during differentiation as observed in our colony formation assay (figure S3A) and demonstrates that scID-seq captures molecular events underlying cellular function. Indeed, other antibodies in our panel also showed quantitative differences between these two cell states in either all, or in a subset of the cells (figure S3C). This highlights the richness and complexity of our dataset and indicates the need to incorporate information on multiple parameters simultaneously in further analyses.

**Figure 2:**
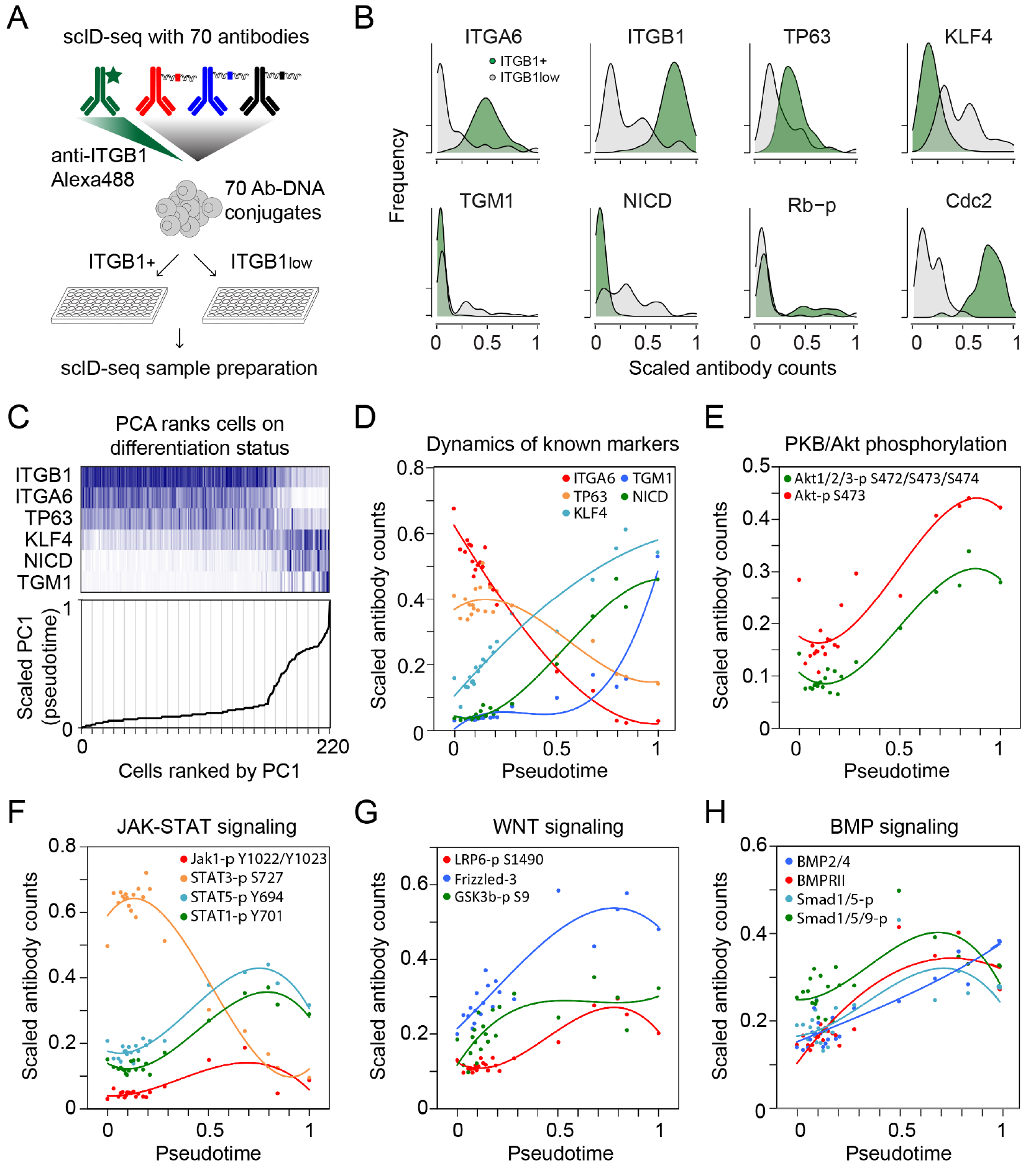
Pseudo-timing reveals dynamic signaling pathway activity over the course of epidermal differentiation. **(A)** Combining scID-seq with FACS-based sorting on ITGB1 levels. Cells were immunostained with fluorescent ITGB1 antibodies in combination with a panel of 70 Ab-DNA conjugates, FACS sorted based on their ITGB1 levels and subjected to scID-seq. (**B)** scID-seq distinguishes ITGB1+ and ITGB1low sorted cells based on known epidermal basal, differentiation and cell-cycle markers. Distributions of normalised and scaled scID-seq counts of selected proteins verified the separation of the ITGB1+ and ITGB1low populations. **(C)** Principal Component Analysis on known markers orders epidermal cells on their renewal and differentiation status. Top panel - markers used for the temporal ordering, color intensity represents scaled antibody counts. Bottom panel - cells were ranked on their (scaled) PC1 loading. Vertical lines indicate the bins used to smoothen the data in subsequent analyses. **(D)** Dynamics of the markers used for PCA, ordered by pseudo-time (scaled PC1 loading) after smoothening. Datapoints indicate bin average and solid lines represent model fit of the data (3rd order polynomial regression). **(E)** Dynamics of two independent phosphorylated Akt/PKB antibodies over pseudo-time. **(F,G,H)** Dynamics of antibodies reflecting the JAK-STAT, WNT and BMP signaling pathways over pseudo-time.

### Pseudo-timing reveals dynamic signaling pathway activity over the course of differentiation

To obtain insight into the progressive changes in signaling pathway activity that occur over the course of epidermal cell differentiation, we aimed to order the cells by their relative differentiation state. We hypothesized that this state can be inferred by examining the combination of expression of known basal and differentiation markers. We performed PCA on selected markers (ITGB1, ITGA6, TP63, NICD, KLF4 and TGM1) with established roles and dynamics during epidermal differentiation, and ranked the cells based on the resulting principal components. This revealed that principal component 1 (PC1; explaining 50% of the variance) recapitulated the expected trajectory of the epidermal differentiation process (figure 2C). We calculated the average of 10 cell bins to smoothen the data, revealing that the basal markers ITGA6 and TP63 are indeed down regulated with distinct kinetics (figure 2D, Kolmogorov-Smirnov (K-S) test, p-value <0.001). Moreover, we observed expected differences in onset and dynamics of induction of the Notch1 Intracellular Domain (NICD), KLF4 and TGM1 during the differentiation process, indicating that the scaled and binned PC1-score reflects the process of epidermal differentiation and can be used as a ‘pseudo-time’ approximation (figure 3D).

**Figure 3:**
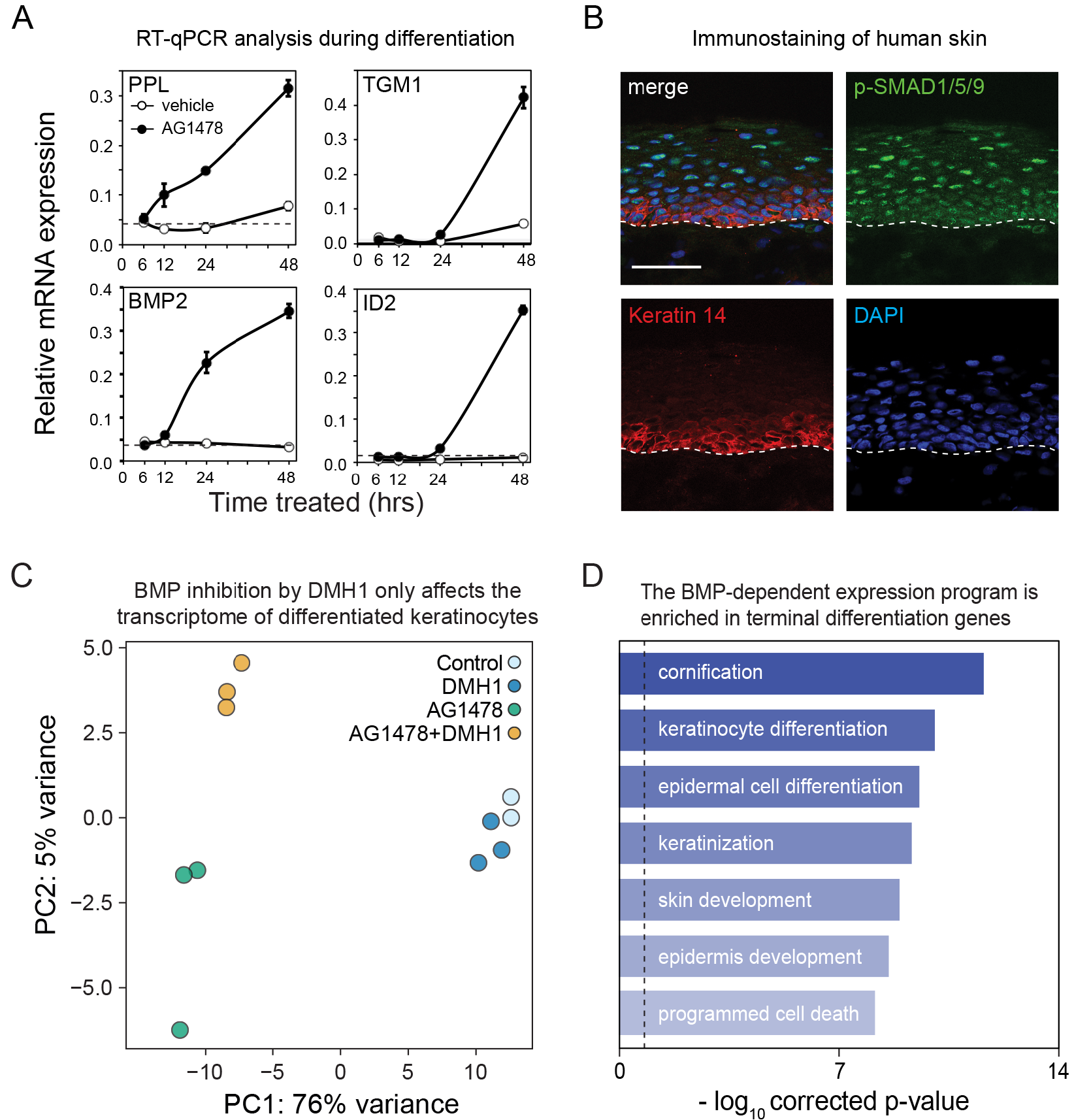
BMP signaling stimulates a terminal epidermal differentiation transcription program. **(A)** BMP2 ligand and its downstream target gene ID2 are activated during in vitro keratinocyte differentiation. Human keratinocytes were induced to differentiate with AG1478 (or DMSO as a control) and samples were harvested at the indicated time-points. RT-qPCR analysis was performed for PPL, TGM1, BMP2 and ID2. **(B)** BMP signaling is activated during epidermal differentiation *in vivo*. Sections of human foreskin were stained with antibodies against phosphorylated SMAD1/5/9. DAPI and K14 antibodies were used to counterstain all nuclei and basal cells, respectively. Scale bar denotes 50 μm. **(C)** Activation of BMP signaling regulates a transcriptional program during epidermal differentiation. Principal component analysis of human keratinocytes induced to differentiate with AG1478 (or the DMSO control) for 96 hours in the presence of the BMP receptor inhibitor DMH1, followed by RNA-sequencing analysis (n=3, except DMSO control n=2). **(D)** BMP dependent genes are involved in late differentiation processes. Top enriched terms from a gene ontology overrepresentation analysis (hypergeometric test, FDR < 0.01)

Next, we mined our data for signaling pathways of which the included antibodies showed concordant effects over pseudo-time and were statistically significantly different between the ITGB1low and the other cells (K-S test, p<0.001, figure S3C). This uncovered several signaling pathways that displayed dynamic behaviour over pseudo-time. For instance, PKB/Akt phosphorylation, as measured by two independent antibodies, was increased upon differentiation as previously described (figure 2E, (Janes et al., 2009)). Besides these expected effects, we found dynamics in 3 additional pathways. The JAK-STAT pathway was activated as evident from increased phosphorylation of JAK1, STAT1 and STAT5, but not the EGFR activated STAT3, during differentiation (figure 2F). In addition, the level of the WNT receptor Frizzled-3 gradually increased with differentiation, as did the activated/phosphorylated form of its co-receptor LRP6 (figure 2G). Moreover, the inactivating serine-9 phosphorylation of GSK3-beta increased and then reached a plateau. This modification helps stabilize cytoplasmic beta-catenin in response to WNT-pathway activation (van Kappel and Maurice, 2017). Finally, the Bone Morphogenetic Protein (BMP) pathway was activated at multiple levels (figure 2H). Our scID-seq panel included antibodies for the BMP2/4 ligand, the type 2 BMP receptor, total SMAD1 levels, as well as two distinct antibodies against phosphorylated SMAD1/5/9. Over pseudo-time we observed increasing levels of the ligand, the receptor, as well as phosphorylated SMADs, reflecting activation of the BMP pathway during differentiation (figure 2H). As our cells are cultured in a defined medium in the absence of feeder cells the signals that activate these pathways must therefore be generated by the cells themselves. Indeed, the increase of the BMP2/4 ligand is consistent with such regulation by the BMP pathway (figure 2H). This suggests that activation of autocrine signaling loops enables cells to coordinate their transcriptional programs and ensure progressive differentiation.

### BMP signaling stimulates a terminal epidermal differentiation transcription program

To validate our findings on the BMP pathway, we induced differentiation of proliferating epidermal cells in culture by inhibiting EGF signaling (Kolev et al., 2008; Mulder et al., 2012). Samples were collected at 6, 12, 24 and 48 hours after the addition of either vehicle (DMSO) or the EGFR inhibitor AG1478. RT-qPCR analysis showed that mRNA expression of the early differentiation marker periplakin (PPL) and the late differentiation marker TGM1 reflected the progression of differentiation over time (figure 3A). In line with our scID-seq results, mRNA expression of the BMP2 ligand was activated upon induction of differentiation (after the 12 hour time-point), whereas the classical BMP-pathway target gene ID2 was induced at a subsequent stage (after 24 hours, figure 3A). Furthermore, the induction of ID2 was dependent on BMP-ligand binding to the BMP receptor, as a recombinant version of the natural BMP-antagonist noggin blocked ID2 expression (figure S4). To investigate BMP pathway activity *in vivo*, we performed immuno-staining with antibodies against phosphorylated SMAD1/5/9 on human skin. The epidermal basal layer was visualised through co-staining with a keratin-14 antibody and we counterstained the DNA of all cells with DAPI (figure 3B). Consistent with our scID-seq and RT-qPCR results, the signal from the p-SMAD1/5/9 antibody was increased in the differentiated (keratin-14 negative) layers of human epidermis, confirming that BMP signaling is activated during differentiation *in vivo*.

These results suggest that autocrine BMP signaling may serve as a positive feed-forward loop to stimulate epidermal differentiation gene expression. To test this hypothesis, we treated cells with AG1478 for 96 hours in the absence or presence of the small molecule BMP receptor inhibitor DMH1 and monitored global gene expression by RNA-sequencing. Vehicle treated cells −/+ DMH1 were included as controls. Principal component analysis indicated that the transcriptomes of non-differentiated and differentiated cells were markedly different (figure 3C, PC1; explaining 76% of the variance). In addition, PC2 (5% of variance) distinguished the DMH1 treated and non-treated differentiated cells, reflecting a transcription program that depends on the BMP pathway (figure 3C). Notably, vehicle control cells were virtually indistinguishable from control cells treated with DMH1, indicating that BMP signaling specifically regulates differentiation, but not proliferation/renewal programs. Moreover, the genes that were dependent on the BMP pathway activity showed highly significant enrichment of genes involved epidermal keratinization and cornification, indicating that autocrine BMP signaling drives transcriptional changes towards terminal differentiation (figure 3D).

### The terminal differentiation transcription factors MAF/MAFB are downstream targets of the BMP pathway

We further explored the mechanistic role the BMP pathway plays in epidermal differentiation by investigating the gene expression program that is influenced by stimulation with recombinant BMPs. First, we treated cells with different BMP ligands and measured the induction of the late differentiation marker transglutaminase I (TGM1) at the protein level. This indicated that most recombinant ligands led to a robust increase of TGM1 (figure S5A). Moreover, simultaneous treatment of cells with the EGFR inhibitor AG1478 and BMP ligands resulted in a synergistic increase of TGM1 protein levels, highlighting opposing roles for these pathways in epidermal biology (figure S5B). Next, we performed RNA-sequencing analysis on cultured human epidermal cells treated with vehicle, AG1478, BMP2/7 or AG1478+BMP2/7 for 48 hours (figure 4A and S6A,B). Comparing the mRNA profiles of these conditions revealed the genes that were responsive to the addition of recombinant BMP2/7 and were also dynamically expressed during AG1478 induced differentiation (figure 4A and S6C). As expected, the top most BMP responsive genes included the ID gene family (figure 4A). Moreover, genes involved in terminal differentiation, including TGM1, MAF, MAFB and ZNF750, displayed BMP induced expression changes, confirming the results obtained with the BMP pathway inhibitor DMH1 (figure 4A,B and 3D). The MAF/MAFB and ZNF750 transcription factor axis drives the epidermal terminal differentiation transcription program (Lopez-Pajares et al., 2015). These results imply that activation of BMP signaling functions upstream of this axis.

**Figure 4:**
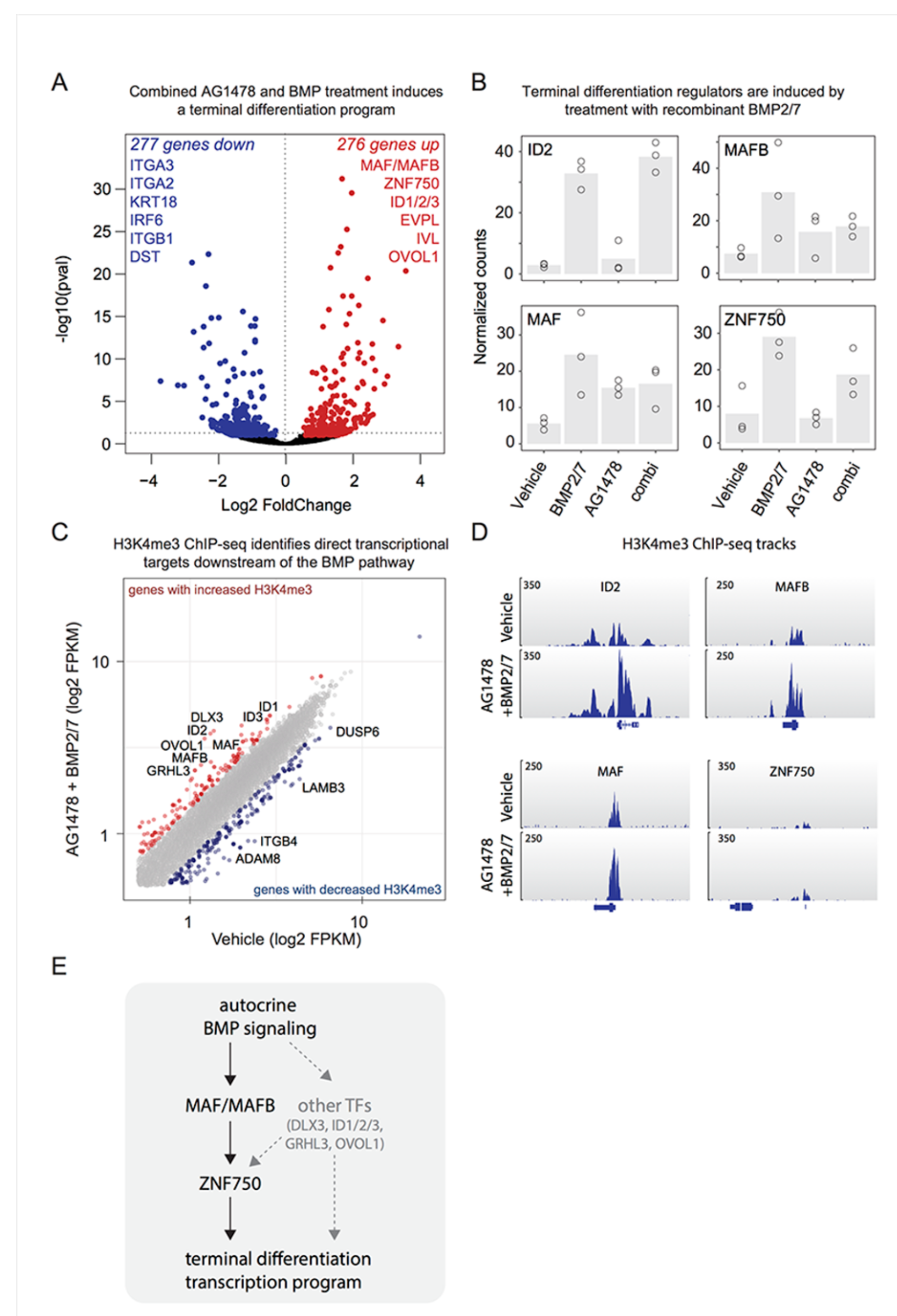
The terminal differentiation transcription factors MAF/MAFB are downstream targets of the BMP pathway. **(A)** Treatment of keratinocytes with recombinant BMPs stimulates epidermal differentiation gene expression. Vulcano plot of cells incubated with AG1478+BMP2/7 (compared to vehicle) for 48 hours, followed by RNA-seq analysis. Vulcano plots of AG1478 and BMP2/7 treatments are presented in figure S6B. Selected differentially expressed differentiation and basal cell markers are indicated **(B)** Recombinant BMPs stimulate expression of transcription factors involved in late epidermal differentiation. **(C)** ChIP-sequencing of the active gene mark H3K4me3 identifies immediate downstream targets after BMP stimulation. Scatter plot of normalised, log-transformed H3K4me3 ChIP-seq signals from cells incubated with and without recombinant BMP2/7 in the context of EGFR inhibition. Genes with 1.5-fold increase or decrease of H3K4me3 signals at their transcription start site are highlighted in red and blue, respectively. Examples of activated and inhibited genes are indicated. **(D)** MAF and MAFB, but not ZNF750, are directly activated by BMP stimulation. H3K4me3 genome-browser tracks of ID2, MAF, MAFB and ZNF750. **(E)** Model of the transcription regulatory network activated by BMP signaling during epidermal differentiation.

To identify transcriptional targets immediately downstream of the BMP pathway, we performed chromatin immunoprecipitation followed by sequencing (ChIP-seq), using H3K4me3 ChIP-seq signals at the transcription start site (TSS) as a proxy for changes in transcriptional activity of a gene. Proliferating epidermal stem cells were treated with vehicle or recombinant BMP2/7 (in combination with AG1478) for 6 hours, after which they were harvested for ChIP-seq analysis. The short duration of the BMP treatment allows us to focus on immediate, rather than secondary effects of BMP pathway activation. This identified 135 genes that showed more than 1.5-fold increase in H3K4me3 signal (fragments per kilobase per million, FPKM) at their TSS, indicating increased transcriptional activity (figure 4C, supplemental table 2). These included the classical BMP targets ID1, 2 and 3, demonstrating that our experimental approach identified direct BMP pathway target genes (figure 4C,D). The terminal differentiation regulating transcription factors GRHL3, MAF and MAFB were among the top set of immediate BMP targets (figure 4C,D). In contrast, the key MAF/MAFB target gene ZNF750 was not directly regulated downstream of the BMP pathway as determined by H3K4me3 ChIP-seq signal (figure 4D). Together, these results place BMP pathway activation immediately upstream of the MAF/MAFB transcription factors. These factors subsequently drive terminal differentiation programs through, among others, ZNF750 (figure 4E, (Lopez-Pajares et al., 2015).

## Discussion

We developed single-cell Immuno-Detection by sequencing (scID-seq) as a highly multiplexed single-cell (phospho-)proteomics approach to study signaling pathway activity in individual human epidermal cells. Measuring 69 (phospho-)proteins per cell demonstrated that, amongst others, Bone Morphogenetic Protein signaling is activated along the differentiation trajectory. Mechanistically, the BMP pathway stimulates the MAF/MAFB/ZNF750-axis to induce a transcriptional program during late stage epidermal differentiation. Previous studies provided indications that stimulation with exogenous BMP ligands increases the expression of cell cycle inhibitory factors and selected differentiation associated genes, suggesting involvement of this pathway in human epidermal differentiation (Botchkarev, 2003; Fessing et al., 2010; Gosselet et al., 2007; Li et al., 2003; Yang et al., 2006). However, it’s timing and its function during the differentiation process remained unclear. Using RNA-sequencing and H3K4me3 ChIP-seq analysis in combination with inhibition and stimulation, we found that BMP signaling activation drives a terminal differentiation transcription program and the MAF/MAFB/ZNF750 transcription factor axis (figures 3 and 4). Our findings have implications for our view on the progressing nature of keratinocyte differentiation. First, regional signaling pathway activity plays crucial roles in patterning and tissue specification during (early) development. Our findings implicate BMP pathway activation as an integral part of the transcription factor network stimulating epidermal differentiation. As the BMP ligand is produced and excreted by the cells into their local environment, our results provide a mechanistic explanation for coordinated expression program progression in a zonated fashion in a tissue context. Second, the identification of the cell intrinsic activation of the BMP pathway through up regulation of its ligand BMP2, in combination with our observation that the BMP pathways is responsible for the stimulation of a specific transcriptional program including late differentiation regulators (e.g. MAF/MAFB, DLX3 and OVOL2), suggests that this pathways is involved in a self-sustaining loop that keeps driving epidermal differentiation forward. Taken together, our results provide insight into the coordination of dynamic transcription programs across cells by incorporating autocrine signaling into cell-intrinsic transcription regulatory networks.

## Materials and methods

### Cell culture and reagents

Primary pooled human epidermal stem cells derived from foreskin were obtained from Lonza. Cells were cultured and expanded as previously reported (Gandarillas and Watt, 1997). Briefly, cells were cultured on a feeder layer of J2-3T3 cells in FAD medium (Ham’s F12 medium/Dulbecco’s modified Eagle medium (DMEM) (1:3) supplemented with 10% batch tested fetal calf serum (FCS) and a cocktail of 0.5 μg/ml of hydrocortisone, 5 μg/ml of insulin, 0.1 nM cholera enterotoxin, and 10 ng/ml of epidermal growth factor) supplemented with Rock inhibitor (Y-27632, 10 μM). J2-3T3 cells were cultured in DMEM containing 10% bovine serum and inactivated with Mitomycin C (SCBT) upon seeding the epidermal stem cells. For experiments epidermal stem cells were transferred to Keratinocyte Serum Free Medium (KSFM) supplemented with 0.2 ng/ml Epidermal Growth Factor and 30 μg/ml bovine pituitary extract from Life Technology until 70% confluent. Cells were treated with AG1478 (10 μM, Calbiochem), DMH-1 (1 μM, RND systems) or BMP2/7 (200 ng/ml, R&D systems). All media were supplemented with 1% penicillin/streptomycin antibiotics.

### Antibody conjugation with dsDNA barcodes

Antibodies and dsDNA were functionalized and conjugated as described (Van Buggenum et al., 2016). Antibody details are provided in supplemental Table 1. In short, antibodies were functionalized with NHS-s-s-PEG4-tetrazine (Jena Bioscience) in a ratio of 1:10 in 50 mM borate buffered Saline pH 8.4 (150 mM NaCl). Then, N3-dsDNA was produced and functionalized with DBCO-PEG12-TCO (Jena Bioscience) in a ratio of 1:25 (oligo list). Finally, purified functionalized antibodies were conjugated to purified functionalized DNA by 4-hour incubation at room temperature in borate buffered saline pH 8.4 in a ratio of 4:1 respectively. The reaction was quenched with an excess of 3,6-diphenyl tetrazine. The conjugation efficiency and quality was checked on an agarose gel, confirming that a substantial amount of DNA conjugated with the antibody. Ultimately, conjugates were equally pooled for staining’s in scID-seq.

### Immuno-staining and single-cell sorting

Cells (> 3 × 10^6^) were harvested with trypsin and cross-linked in suspension by incubating for 10 minutes with 4% paraformaldehyde (PFA) in PBS following a quenching step of 5 minutes with 125 mM Glycine in PBS. Removal of PFA and Glycine occurred through washing twice with wash buffer (0.1× Pierce™ Protein-Free Blocking Buffer from Thermo in PBS). Then, cells were blocked in 500μl blocking buffer (0.5× 0.1× Pierce™ Protein-Free Blocking Buffer, 200 μg/ml boiled salmon sperm DNA, 0.1% Triton-X 100, in PBS) at room temperature for 30-60 min. Staining with the conjugate mix occurred overnight at 4°C in 500 μl blocking buffer and pre-staining’s were performed at room temperature for 1-2 hours. After each staining, cells were washed 3x in 5ml wash buffer. Cells were sorted single cell with the BD FACSAria SORP flow cytometer (BD biosciences) in 96 well PCR plates containing 1μl release buffer (10 mM DTT in 15mM Tris, pH 8.8) and 7ul Vapor-lock (Qiagen). For selection of ITGB1 negative cells, a primary and secondary pre-stain was done with 2.5 μg/ml anti-ITGB1 (P4D1) and 1:1000 Alexa488 goat anti-mouse (Life technology, 1484573). Plates were stored at −20°C until use.

### Barcoding and library preparation for next generation sequencing

For the library preparation 3 PCR steps were performed to amplify the antibody barcodes and to add barcodes specific for the well and the plate of each cell. The barcoding occurred with the same sequences used in ID-seq (Buggenum et al., 2017). For the first PCR step 15 cycles were run after adding to each well a 4 ul reaction mix containing the Herculase II Fusion DNA Polymerase (Agilent), dNTPs, 5x Herculase buffer and 0.1μM amplification primers (Forward 5′-CACGACGCTCTTCCGATCT-3′, Reverse 5′-TCGCTTATCTGTTGACTGAT-3′). Directly after the first PCR step, 5 extra cycles were run after adding 1 ul mix containing Herculase buffer 0.2 μM forward amplification primer and 0.2 μM reverse well barcoding primer. Then all material was pooled per plate, Vapor-lock was removed and a clean-up was performed with the QIAquick PCR Purification Kit, an EXO1 treatment to degrade remaining primers followed by another purification. Another 5 cycles were run in PCR 3 with a 20 ul reaction containing pooled and purified plate sample and 0.1 μM plate barcoding primers (Fw_long and specific plate reverse). After repeating the clean-up, the libraries were checked on agarose gel and with the Bioanalyzer (Agilent) to confirm the size of the DNA fragments (expected size around 185 bp).

### Data analysis

Sequence data from the NextSeq500 (Illumina) was demultiplexed using bcl2fastq software (Illumina). Then, all reads were processed using our dedicated R-package (Buggenum et al., 2017). In short, the sequencing reads were split using a common “anchor sequence” identifying the position of the UMI sequence, Barcode 1 (antibody specific) and Barcode 2 (well specific) sequence. After removing all duplicate reads, the number of UMI sequences were counted per barcode 1 and 2. Finally, barcode 1 (“antibody)” and barcode 2 (‘well’) sequences were matched to the corresponding. For scID-seq, a threshold was set based on the total UMI count per well difference between qualitative cells and low quality cells or empty wells. Cell populations pre-stained with specific antibody-barcodes were indexed via determining the enriched pre-stain barcode per cell. Per experiment, cells were normalized through subsampling. Principle component analysis was done with the R-package “pcaMethods”. Additional statistical analysis and visualization of the data was done in R and Excel.

### Colony formation assay

Epidermal stem cells were sparsely seeded on a feeder layer in a 6 well plate (500 cells/well) and cultured for at least 7 days to form colonies. After colony formation, cells were washed with PBS and fixed by incubating 10 minutes with 4% paraformaldehyde (PFA) in PBS. PFA removal occurred with 3 PBS washes. For the imaging of the colonies, a DNA stain was performed with DRAQ5 (1:4000, Biostatus) for 1 hour at room temperature. After 3 PBS washes, plates were scanned with the Odyssey system and the number of colonies quantified.

### Immuno staining of human skin

Frozen sections of human foreskin were a kind gift from Prof. Fiona Watt and were obtained with informed consent and appropriate ethical review. Sections were fixed for 15 minutes with 4% paraformaldehyde at RT followed by 3 washes with PBS and permeabilisation with 0.2% triton-X100 in PBS for 10 minutes at RT. After blocking with 10% Bovine Serum in PBS for 1 hour, the sections were stained with antibodies against phosphorylated-SMAD1/5/9 (41D10, Cell Signaling) at 1:200 dilution in blocking buffer overnight at 4 degrees Celsius. Sections were washes 3x with PBS and stained with secondary antibodies (1:2000 dilution) and DAPI (1:5000) for 90 minutes at RT. Images were acquired using a Leica IR laser confocal microscope.

### RT-qPCR

Isolated RNA (Qucki-RNA TM MicroPrep, Zymo Research) was used for quantitative PCR analysis using iQTM SYBR Green Supermix with 20μM reaction volume, scanned on CFX-96 machine. Per gene, −2^Ct values were calculated and normalized to 18S RNA levels.

### Transcriptome analysis with CEL-seq2

Isolated RNA (Qucki-RNA TM MicroPrep, Zymo Research) was used for transcriptome analysis via a slightly modified CEL-seq2 procedure (Hashimshony et al., 2016). See table 1 primer sequences. In short, 100 pg purified RNA was used in 2 μl reverse transcription reactions containing Maxima H minus reverse transcriptase (ThermoFisher). The reactions were covered with Vapor-Lock (7 μl, Qiagen). Different primer sequences were designed and used (Supplementary table 1), allowing 63 nt long read 1 of mRNA, and 14 nt long read 2 with the sample barcode and UMI. NextSeq500 (Illumina) was used for sequencing.

**Table 1.**
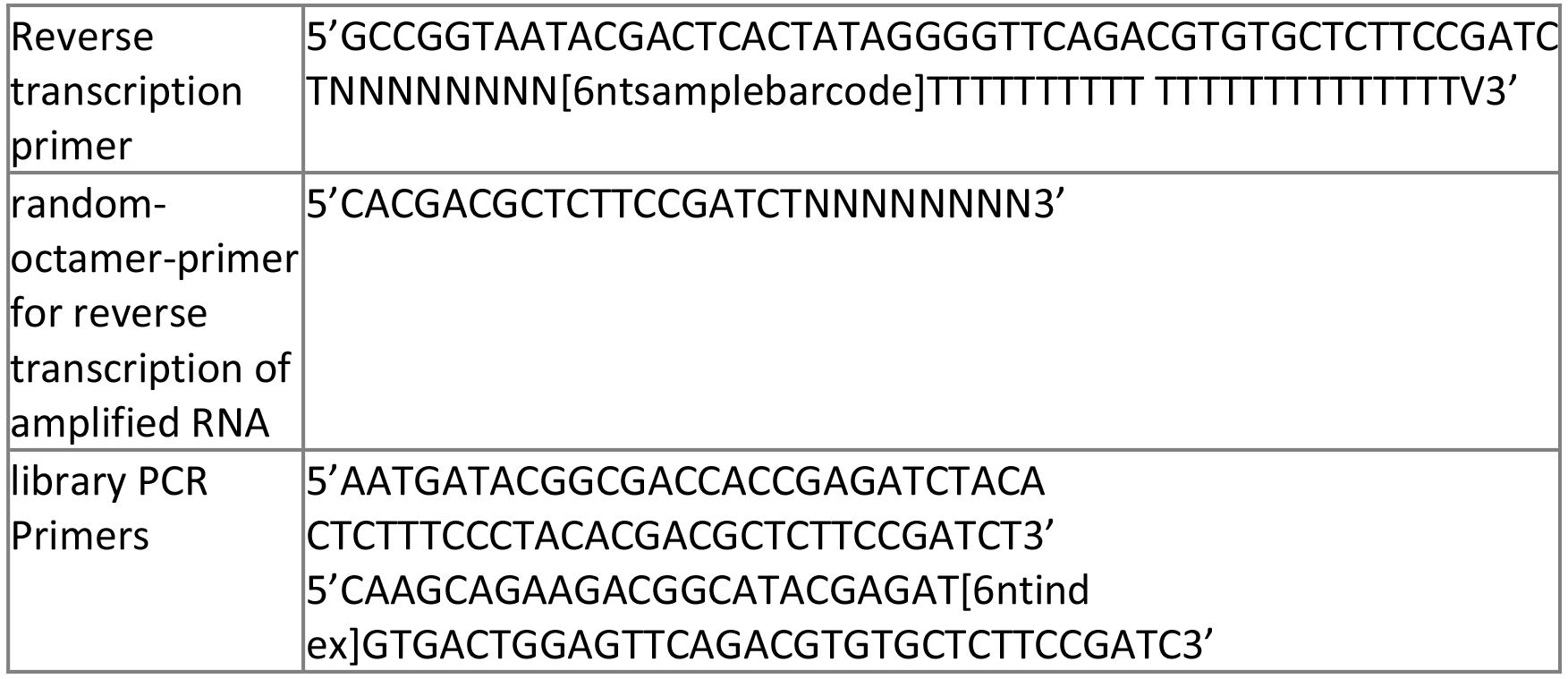
Primer sequences used during CEL-seq2 procedure.

### ChIP-sequencing

Cells in KSFM (2.5×10^6^) were incubated with vehicle/DMSO or BMP2/7+AG1478 for 6 hours. After harvesting by trypsinisation, the cells were cross-linked with 1% formaldehyde in PBS for 10 minutes, quenched for 5 minutes with 125 mM glycine in PBS and washed in PBS at 4 degrees Celsius. Cross-linked cells were incubated 4 hours in lysis buffer (5 mM Tris-HCl pH 8.0, 85mM NaCl, 0.5% NP40, 1X PIC) on ice, following 50 strokes with a dounce homogenizer to enrich for nuclei. Nuclei were sonicated in sonication buffer (50 mM Tris-HCl pH 8.0, 10 mM EDTA, 0.1% SDS, 150mM NaCl and 0.5% deoxycholic acid) to get an average chromatin fragment length of 500 bp. Chromatin extracts were incubated with 1 μg H3K4me3 antibody (ab8580, AbCam) overnight. Antibodies were captured for 4 hours with 100 μl protein G-coated magnetic beads (Life Technology). Subsequently, beads were washed 5X in RIPA buffer. The retrieved chromatin was reverse cross-linked overnight at 65°C followed by a 1 hour incubation step with proteinase K (1μg/μl) and RNAse A (1μg/μl) at 37°C. The DNA was purified with the Qiaguick PCR purification kit. Using the KAPA Biosystems kit (#KK8504), between 0.5 and 5ng ChIP-derived DNA was trimmed, A-tailed, provided with Nextflex adaptors and amplified with 10 PCR cycles. Subsequently, DNA was size-selected with the E-Gel^®^ iBase™ Power System (Invitrogen) to purify for fragments between 300 and 400bp. The libraries were quantified on the Agilent 2100 Bioanalyzer and evaluated by qPCR to confirm representation of enrichments at specific loci. The libraries were sequenced with the Nextseq500. Reads were quality checked and aligned to the human hg19 genome with the Burrows-Wheeler Alignment tool (BWA) and processed with SAMtools to generate BAM files. Peaks were called from BAM files using “Model-based analysis of ChIP-Seq version 1.4” (MACS14) with a p-value cut-off of 1×10^−8^.

### Data analysis and availability

CEL-seq2 sequencing data was processed using as described (Hashimshony et al., 2016). In brief, high quality reads were filtered, and used for mapping. QC analysis of the dataset is described in detail in R-notebook file available from https://github.com/jessievb. In brief, the count matrix was loaded as Seurat object, and used to visualize number of total UMI counts, number of genes and number of mitochondrial genes per sample. One out of three vehicle controls in the DMH-1 experiment did not pass the QC checks (with 50% less reads/sample) and was removed from further analyses. Data analysis and visualization is described in detail (https://github.com/jessievb). In brief, the count matrix was loaded into DESEq2 dataobject, to allow easy normalization, filtering, PCA and differential expression analysis. For analysis of the CEL-seq data, the DESeq2 R-package (Love et al., 2014) was used to normalize UMI counts, perform PCA analysis and determine differential expressed genes. Sequencing data are available from GEO under series number GSE115926.

## Author contributions

RvE, JvB and ST designed and performed experiments, analysed and interpreted data and wrote the manuscript. JH performed experiments. KM conceived the project, designed experiments, interpreted data and wrote the manuscript.

## Acknowledgements

We thank members of the Mulder lab for fruitful discussion and assistance and Dr. H Zhou and C. Albers for critical reading of the manuscript. This work was supported by a VIDI grant from the Netherlands Organisation for Scientific Research (NWO-VIDI) to K.M.

---

**Figure S1:**
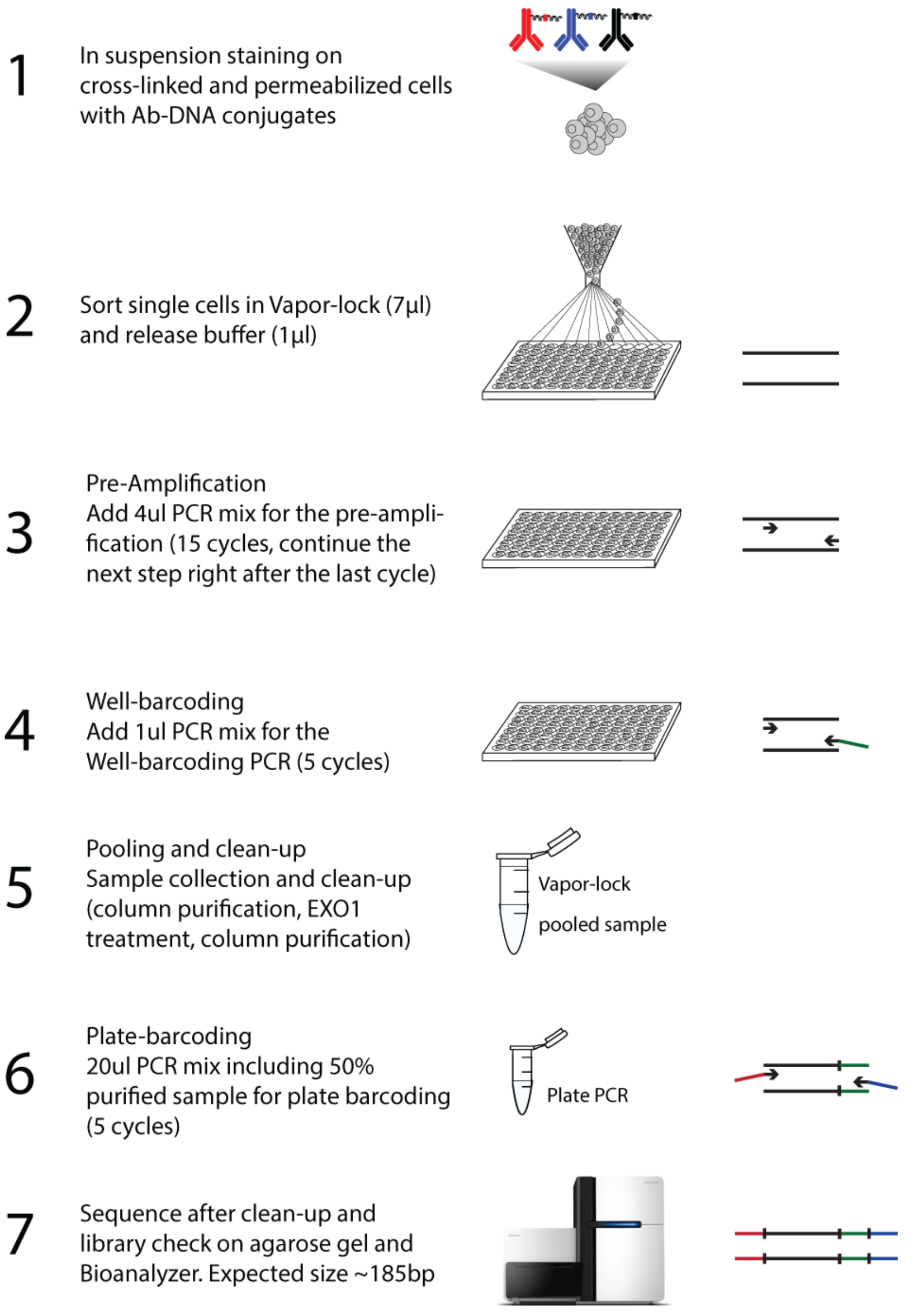
Schematic representation of the single-cell ID-seq work-flow. The scID-seq procedure entails two crucial modifications compared to the original ID-seq protocol. First, cells are stained with the antibody-DNA conjugates in suspension, to allow single-cell sorting. Second, a pre-amplification step increased the yield and complexity of the single-cell libraries. This pre-amplification is done prior to adding a cell-specific barcode to the PCR products. From this point on, the library preparation is the same as described for ID-seq.

**Figure S2:**
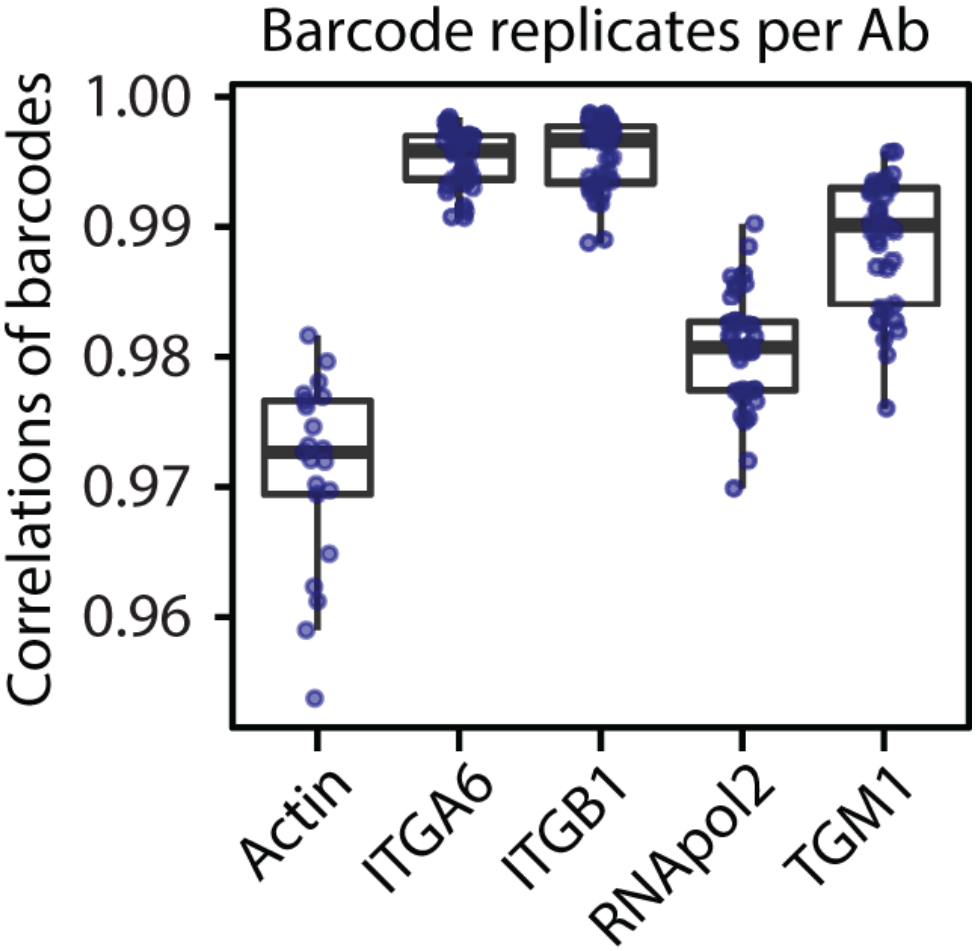
Nine technical replicates per individual cell highlight the reproducibility of scID-seq. Cells were stained with a mixture of 45 antibody-DNA conjugates consisting of 5 different antibodies that were each separately conjugated to 9 independent DNA-barcodes. The Pearson correlation among the 9 measurements for each antibody across individual cells indicates the reproducibility of scID-seq to quantify relative protein levels.

**Figure S3:**
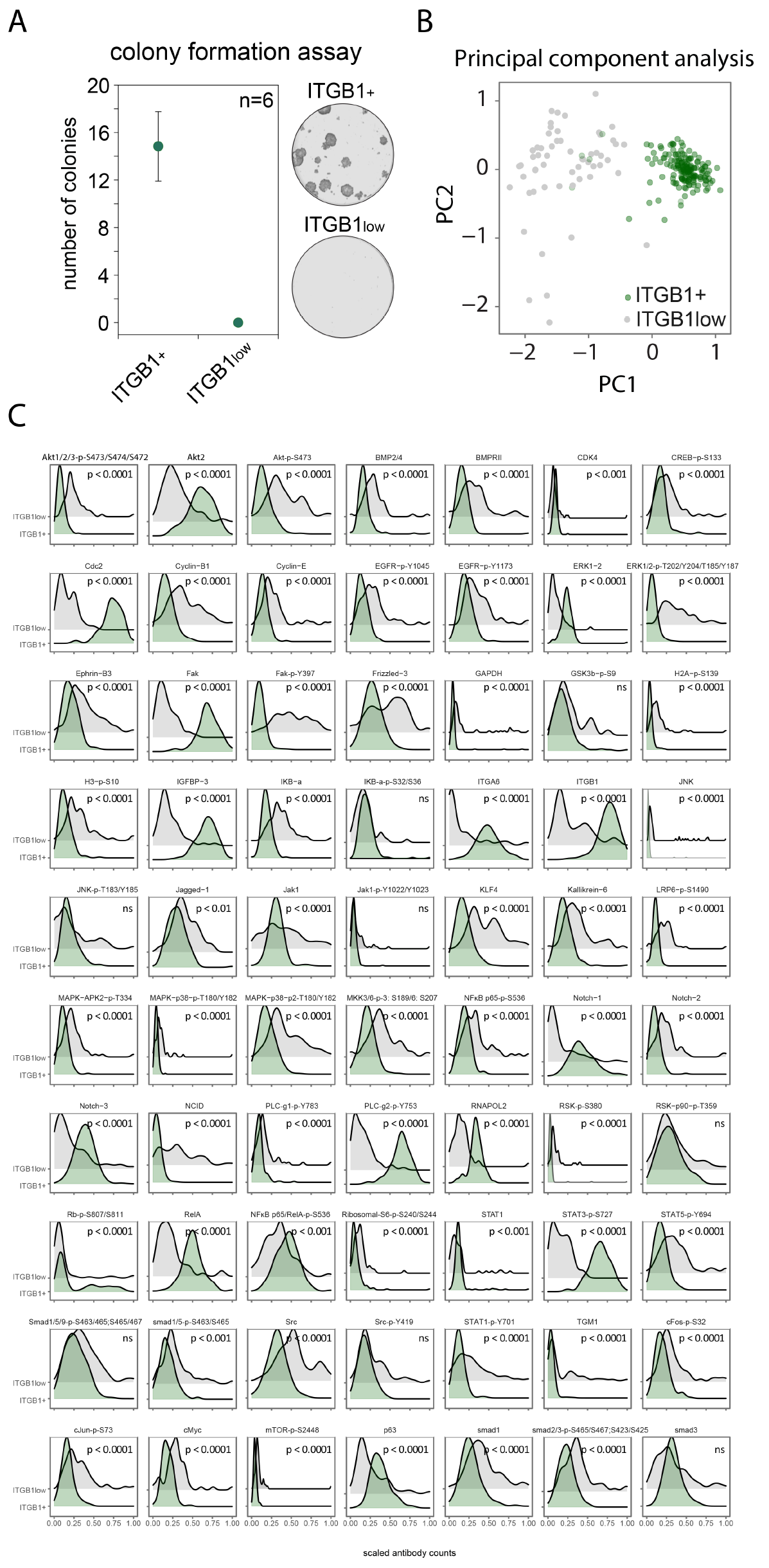
scID-seq on cells selected on ITGB1 level by FACS. **(A)** Cells with low ITGB1 surface levels represent differentiated keratinocytes. Cultured human keratinocytes were isolated based on their ITGB1 surface expression by FACS and subjected to colony formation assays. **(B)** ITGB1+ and ITGB1low cells display distinct scID-seq profiles. Human keratinocytes were subjected to staining with a panel of 70 antibody-DNA conjugates, isolated based on their ITGB1 surface level by FACS and analyzed with scID-seq. Principal component analysis separated the ITGB1+ and ITGB1low populations. **(C)** Individual antibodies display dynamics between ITGB1+ and ITGB1low cell populations. Distributions of scaled signals of ITGB1+ and ITGB1low populations for 70 antibody-DNA conjugates. Statistically different distributions are indicated (Kolmogorov-Smirnov test, p<0.001).

**Figure S4:**
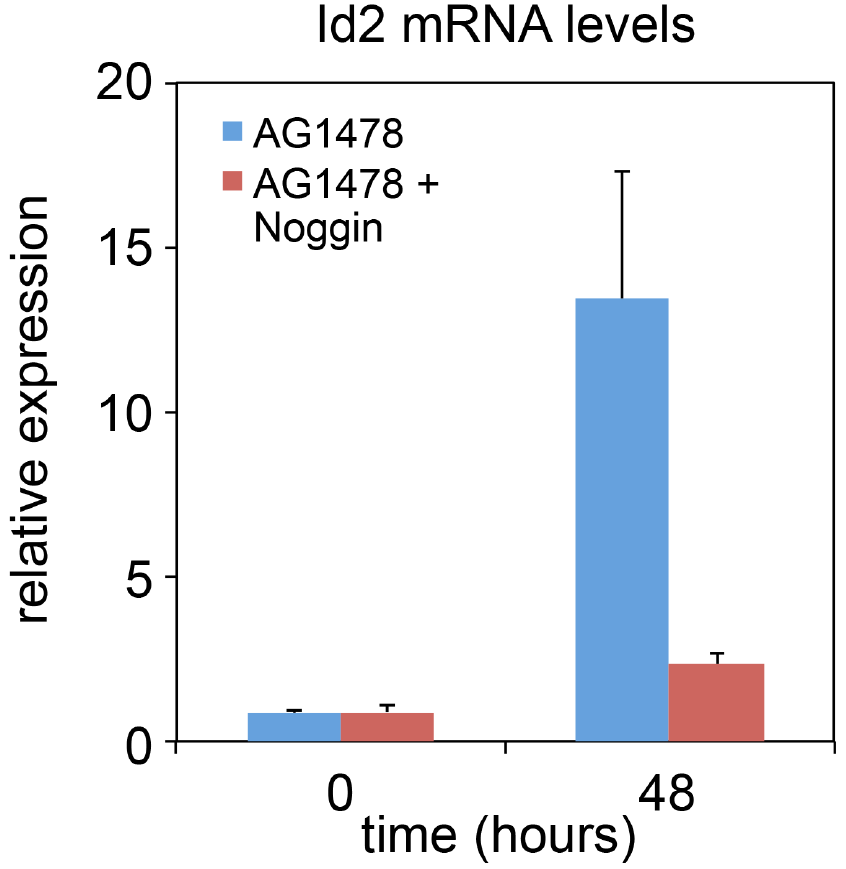
The BMP pathway target gene ID2 is induced during differentiation in a BMP-BMPR interaction dependent manner. Human keratinocytes were induced to differentiate with the EGFR inhibitor AG1478 for 48 hours and subjected to RT-qPCR analysis of the ID2 gene.

**Figure S5:**
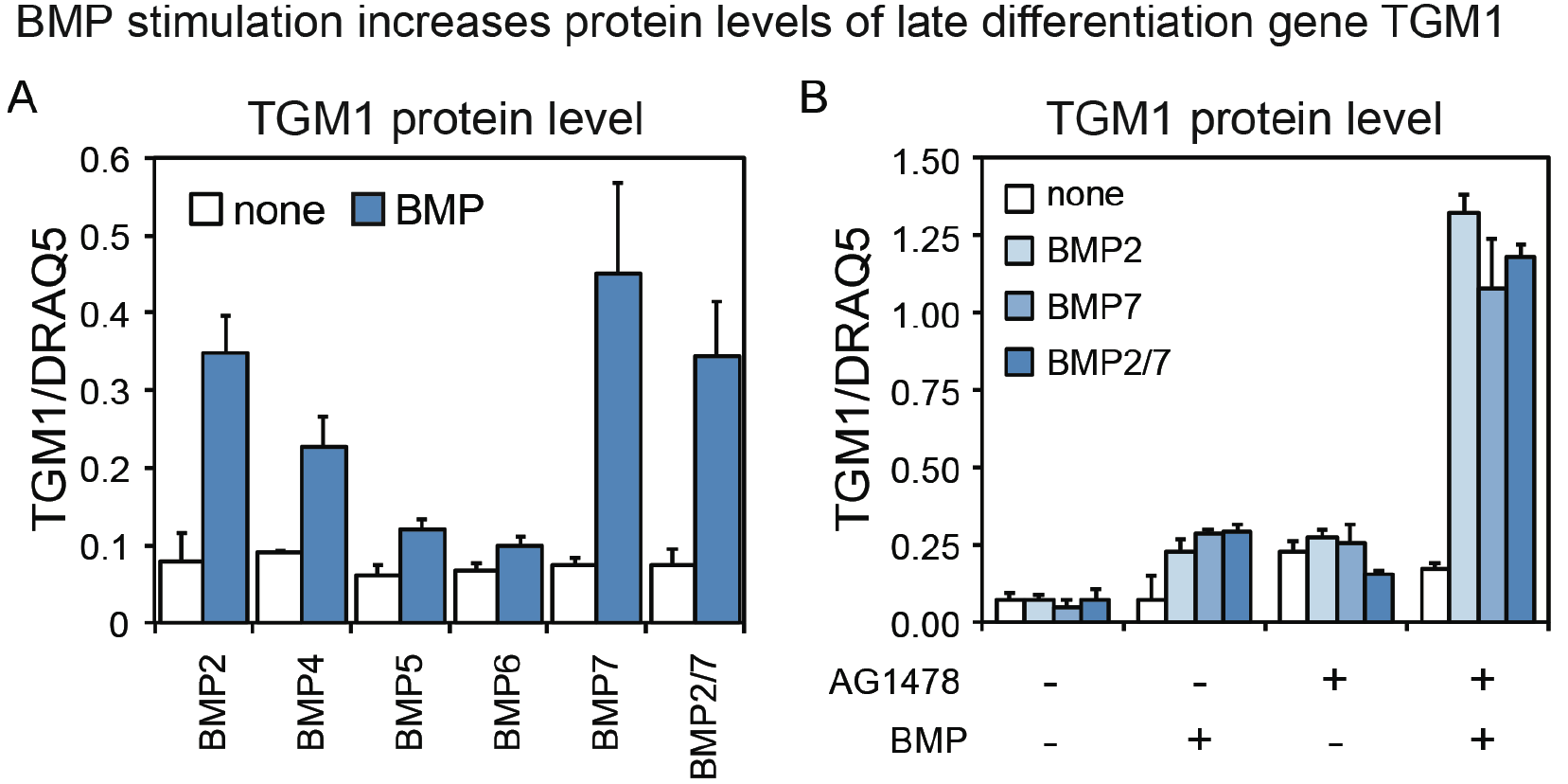
Stimulation with recombinant BMPs induces TGM1 protein levels alone and in synergy with EGFR inhibition. **(A)** Recombinant BMPs stimulate endogenous TGM1 protein expression. Cells were stimulated with the indicated recombinant BMP proteins for 48 hours and subjected to In-Cell-Western analysis using TGM1 specific antibodies. Measurements were corrected for cell density using the DNA staining agent DRAQ5. n=3 −/+ SD **(B)** Recombinant BMPs and AG1478 syngergise to stimulate endogenous TGM1 protein expression. Cells were stimulated with the indicated recombinant BMP proteins for 48 hours in combination with the EGFR inhibitor AG1478 and subjected to In-Cell-Western analysis using TGM1 specific antibodies. Measurements were corrected for cell density using the DNA staining agent DRAQ5. n=3 −/+ SD

**Figure S6:**
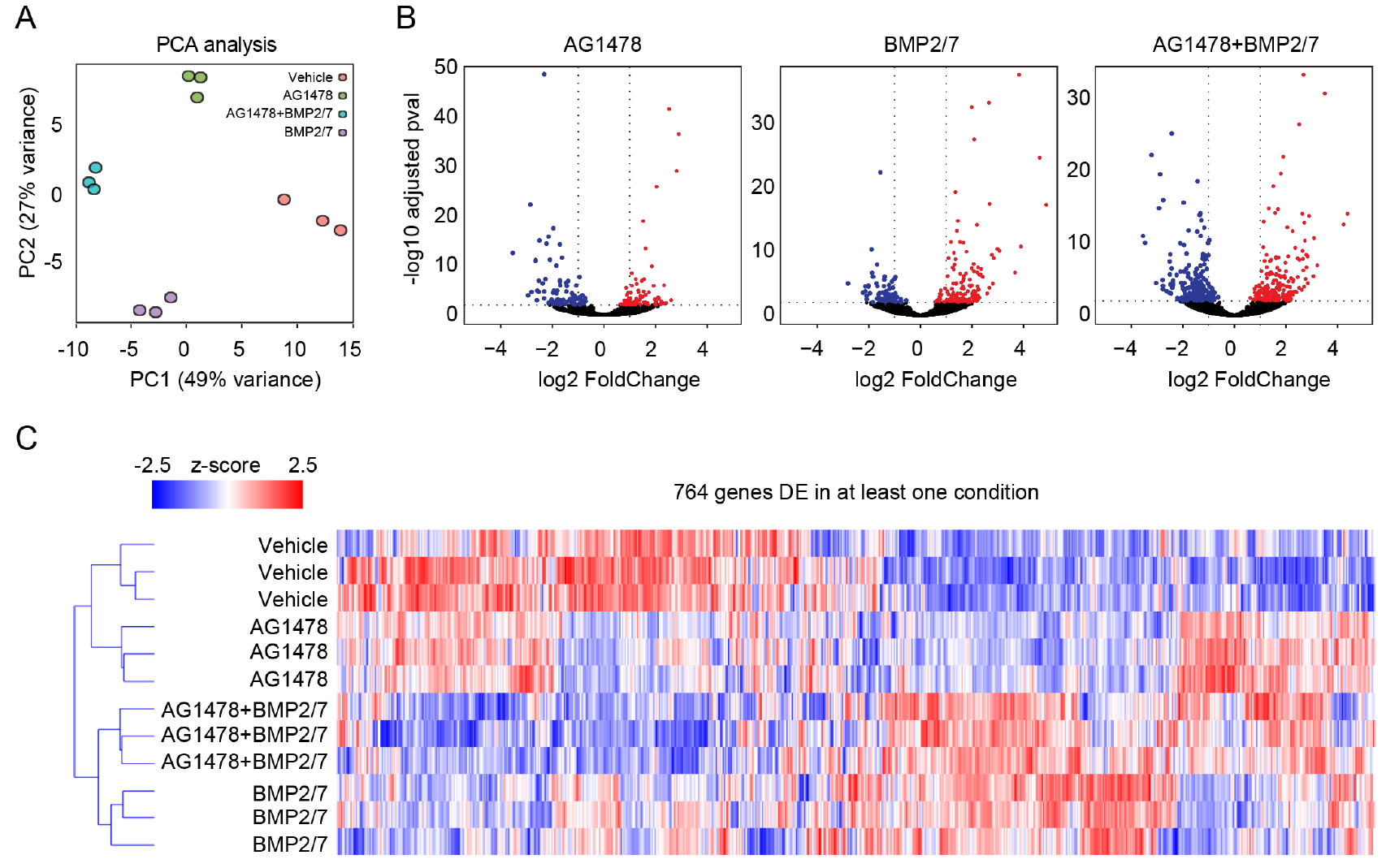
RNA-sequencing analysis reveals BMP responsive expression programs in human keratinocytes. **(A)** Recombinant BMPs, AG1478 and the combined treatment lead to distinct transcriptional responses. Cells were stimulated with the indicated treatments for 48 hours and subjected to RNA-sequencing. Principal component analysis of differentially expressed genes indicates condition specific transcriptional effects. **(B)** Identification of differentially expressed genes per condition. Volcano plots indicate the log2 fold-change of mRNA expression between vehicle and the indicated condition on the x-axis. The y-axis represents the-log10 transformed p-value (FDR corrected t-test). **(C)** Hierarchical clustering highlights condition specific differences in transcriptional responses. Heatmap of differentially expressed genes (z-score normalised across the samples) clustered on Pearson’s correlation with average linkage for both genes and samples.

